# The effectiveness of clove oil and cautery disbudding methods on preventing horn growth in dairy goat kids

**DOI:** 10.1101/325159

**Authors:** Melissa N Hempstead, Joseph R. Waas, Mairi Stewart, Vanessa M. Cave, Amanda R. Turner, Mhairi A. Sutherland

## Abstract

The effectiveness of clove oil and cautery disbudding on horn growth was evaluated in goat kids. The study used 243 Saanen doe kids (4±1.0 days old; mean ± SD) on two commercial dairy goat farms, and were disbudded with either (i) clove oil injection (CLOVE), (ii) a cautery iron and bud removed (BUDOFF), or (iii) a cautery iron with bud left intact (BUDON). Each kid received a different treatment per bud, which were balanced between buds (left and right) and randomly allocated. A trained observer monitored bud growth following treatment for 3 months recording either: N: no growth, H: normal horn, S: abnormal horn (scur), or SC: soft, fibrous lump (scorn). After the final observation, buds were assessed for the probability of detecting (i) success (no growth), (ii) scurs, (iii) horns or (iv) scorns [with 95% CI]. The probability of success for BUDOFF (0.77 [0.63, 0.87]) was higher than for BUDON (0.20 [0.11, 0.34]) and CLOVE (0.09 [0.04, 0.18]; *P* ≤ 0.05). Furthermore, the probability of success for BUDON was higher than for CLOVE (*P* ≤ 0.05). The probability of scurs was higher for CLOVE (0.72 [0.63, 0.80]) than BUDOFF (0.25 [0.17, 0.34]) and BUDON (0.30 [0.21, 0.39]; *P* ≤ 0.05). There was no difference in the probability of scurs for BUDOFF and BUDON (*P* > 0.05). The probability of horns was higher for CLOVE (0.21 [0.15, 0.29]) than BUDON (0.02 [0.01, 0.06]; *P* ≤ 0.05); horns were not observed for BUDOFF. The probability of scorns for BUDON, the only treatment that led to scorns, was 0.41 (0.25, 0.60). These results suggest that BUDOFF was more effective at preventing growth than CLOVE and BUDON and appears the most effective method, of the methods tested, for disbudding kids. Future research should explore different clove oil administration methods or other alternatives to cautery disbudding that may be both efficacious and cause less pain.

## Introduction

Dairy goat kids are routinely disbudded, usually within the first week of life [1]; the practice involves the destruction of the horn buds to prevent horn growth. The predominant method involves using a hot cautery iron to cauterise and remove horn buds. Disbudding is carried out to reduce the risk of injury to other goats [2] and their human handlers, and allows for more space at the feed rail or in lying areas [3]. Current disbudding methods for goat kids have been adapted from those used for calves, such as cautery disbudding, caustic agents and surgical methods [4–6].

It is generally accepted that cautery disbudding causes pain and distress in goat kids as evidenced by intense and frequent vocalisations, leg shakes during the procedure [7], elevated plasma cortisol concentrations and increased frequencies of head shaking, rubbing and scratching post-disbudding [8–10]. Furthermore, potential complications associated with cautery disbudding include second-or third-degree burns, inflammation, thermal injury to the skull and brain (causing necrosis), infection and an increased risk of mortality [11–13].

If disbudding of goat kids is not performed, then adult goats can be dehorned; this involves surgical removal of the horns, which creates an opening into the frontal sinus and causes more pain than disbudding [5]. It can also lead to complications such as prolonged healing, discharge/infection, inflammation, regrowth of horns, dehiscence or even death [14,15]. Therefore, disbudding is the preferred method for preventing horn growth.

There is limited research on the efficacy of different disbudding methods (including cautery disbudding) on horn or scur growth in goat kids or calves. Disbudding is considered successful if no horns or scurs grow (i.e. all horn bud tissue is destroyed). Scurs can be defined as distorted horn regrowth following disbudding [15]. Problems associated with scurs include being aesthetically unpleasing, breaking off easily causing open wounds and increasing the risk of infection and potential abnormal growth back towards the animal’s head, requiring surgical removal. Cautery disbudding can be performed by either (i) totally removing the ring of tissue containing the horn bud cells or (ii) cutting/burning a circular ring around the horn bud but leaving it intact. It is unclear which method of cautery disbudding is most effective in preventing scurs or horns.

A recent study [9] used the physiological and behavioural responses of dairy goat kids to evaluate alternatives to cautery disbudding (i.e. caustic paste and cryosurgical disbudding, and clove oil injection). Clove oil injection elicited a similar cortisol response and number of head shakes and scratches as cautery disbudding, indicating a similar experience of pain [9]. Even though the pain response to clove oil injection appeared to be similar to that generated by the cautery iron, the clove oil method caused less tissue damage [9]. Consequently, clove oil injection may result in faster healing times and lower rates of infection or skull or brain injury. Caustic paste and cryosurgical disbudding appeared to cause more pain than cautery disbudding [9], and therefore were not included as treatments in the present study.

Clove oil, which was traditionally used in dentistry as a topical analgesic and antiseptic [16], contains a high concentration of eugenol (i.e. 80-85%), which has been shown to cause cellular necrosis (and inflammation) in the oral mucosa of rats [17]; it can also be highly cytotoxic for human skin cells [18]. Recently, clove oil has been shown to cause local cellular necrosis of horn bud tissue resulting in arrested horn growth in calves [19] and goat kids [20]. Further research is required to evaluate the effect of clove oil on horn and scur growth in goat kids.

The objective of this study was to evaluate the effectiveness of clove oil injection and two methods of cautery disbudding (i.e. horn buds removed vs. left intact) on horn bud growth in dairy goat kids. We predicted that clove oil injection would result in similar levels of horn and scur growth as cautery disbudding, based on the success rates (100%) reported by others [20]. We also predicted that cautery disbudding with horn buds removed, would result in less horn and scur growth than leaving the horn buds intact due to increased potential for complete cell destruction.

## Materials and methods

The Ruakura Animal Ethics Committee approved the use of animals prior to the commencement of the study (Protocol No. 14213).

## Pilot study

A pilot experiment was carried out to ensure that administration of clove oil into the horn bud had no detrimental effect on the skull and brain of goat kids before further studies were conducted. On a commercial dairy goat farm in the Waikato region of New Zealand, 10 Saanen doe kids were selected from unwanted stock based on age (i.e. 2-3 days old) in June 2017. The animals were injected with 0.2 mL of clove oil into each horn bud using the procedure described in Hempstead et al. [9]. Kids were reared in a small barn separate from the farmer’s replacement stock and fed using a 10 L bucket feeder with 6 teats. Two weeks after clove oil injection, 5 kids were euthanized by a veterinarian. The remaining 5 kids were euthanized 1 week later (i.e. 3 weeks after clove oil injection) to assess the effects of clove oil over time. After euthanasia, the bodies were transported to a post mortem facility at the Ruakura Research Centre, in Hamilton, New Zealand, and gross examination was then performed by a veterinary pathologist.

Firstly, the skin over the head was visually assessed and then the skin was removed so that the outside of the skull could be examined. Next, the head was cut transversely, just caudal to the horn buds using a commercial meat band saw. The brain was removed from the front part of the skull and the inner surface was examined for evidence of damage (e.g., perforation, hyperaemia) or inflammation beneath the horn bud sites. The dorsal surfaces of cerebral hemispheres beneath these sites were examined for ulcerations.

Large black scabs covered the horn buds 2 weeks post-injection of clove oil. There were localised dark patches on the skull below the horn buds for the 5 kids euthanized 2 weeks after treatment, with no evidence of damage on the inside of the skull or the brain. At 3 weeks post-injection, patches of newly healed skin were observed as well as scurs in 3 out of the 5 kids. Discolouration of the skull beneath the horn buds was apparent in only one kid. There was no evidence of inflammation, perforation or infection in the skull, meninges or brain associated with the clove oil injection for any of the kids.

## Animals and housing

This study was conducted on two private commercial dairy goat farms within the Waikato region in New Zealand between June and December 2017. A total of 243 Saanen doe kids (4 ± 1.0 days old; mean ± SD) were used (Farm A: 189 kids; Farm B: 54 kids) and were selected for inclusion based on age (2–6 days old) and size of horn buds (< 16 mm). Kids had an average body weight of 4.0 ± 0.55 kg (range: 2.7 – 6.1 kg). Kids were housed in pens (3.5 x 2.0 m) with approximately 15 kids/pen until they were 2 weeks old or were feeding independently, at which point they were moved to larger pens (9.0 x 5.0 m) with approximately 45 kids/pen.

The kids were reared as per routine practice on each farm [21]. The ground within the pens was covered with untreated pine shavings (approximately 10 cm deep). Each pen had *ad libitum* access to milk replacer (SprayFo, AgriVantage, Hamilton, New Zealand; mixed according to packet instructions) in feeders (Milk Bar, Waipu, New Zealand) with 10 teats/feeder (36 L capacity) and fresh water in a trough.

## Experimental design

A power analysis for a binary outcome bio-equivalence trial was conducted based on 80% power, a 5% significance level and an equivalence range of +/− 0.15. We used a randomized spilt-plot design with a different treatment per horn bud, with treatment day, kid, and farm as blocking variables. A replicate consisted of all pairs of treatments and whole replicates were completed per treatment day. Horn buds were randomly allocated treatments (n = 162 horn buds/treatment) balanced for kid age. Kids were given a coloured collar for treatment identification within the pens as they were housed with others not experiencing any treatment. The same veterinarian performed all treatments. The experiment was conducted on 8 treatment days over a 2.5-week period. Kids were fed approximately an hour before treatment and then taken (one at a time) from their home pen and placed in a restraint device (described in Hempstead et al. [10]). Treatments were performed in an alleyway alongside the home pens and treatment order was randomly assigned. Hair covering the horn buds was removed with an electric clipper (Laube, 505 cordless kit, Shoof, Cambridge, New Zealand) to expose the horn buds. Prior to treatment, kids received an oral non-steroidal anti-inflammatory drug (Loxicom 0.5 mg/mL oral suspension for dogs, Norbrook Laboratories Ltd, Newry, UK; 0.2 mg/kg BW) and a cornual nerve block using lidocaine (Lopaine 2%, 20 mg/mL, Ethical Agents, Auckland, New Zealand; 0.1 mL/horn bud) to reduce pain associated with treatment.

Treatments included:

i. BUDOFF: Disbudding using a cautery iron (“Quality” electric debudder, 230 V, 190 W; Lister GmbH, Lüdenscheid, Germany), which was heated for 20 min (to reach c. 600°C) prior to being pressed to each horn bud for a total of 5.9 ± 1.09 s (mean ± SD). Horn buds were then removed by pressing the iron down and rotating so the skin was cut and the buds forcibly flicked out, as described in Hempstead et al. [8].
ii. BUDON: The same procedure as for BUDOFF, except that the horn buds were cut but not removed. The iron was held to each horn bud for a total of 4.8 ± 1.08 s on average.
iii. CLOVE: Clove oil (C8392, 100mL, 83-85% eugenol, Sigma-Aldrich, Saint Louis, MO) was injected (0.2 mL; [20]) laterally into the centre of each horn bud at a 45° angle between the ear and muzzle (20.9 ± 8.39 s; mean completion time ± SD); details of the procedure are described in Hempstead et al. [9].

After treatment, BUDON and BUDOFF wounds were sprayed with antibacterial spray (Tetravet, Bayer New Zealand Ltd., Auckland, New Zealand) to prevent infection. Horn buds treated with CLOVE did not receive spray as there were no open wounds. Kids were then returned to their home pen and were monitored for 2 h post-treatment to ensure no complications associated with treatments occurred. The health status and horn bud growth of the goat kids was assessed for 5 months post-treatment. Any kids that died over the course of the experiment were examined post-mortem by a veterinarian to determine cause of death.

## Horn bud growth categories

Horn bud growth categories were defined before the start of the experiment. Each horn bud was categorised based on whether it displayed normal horn growth (H), abnormal growth or scurs (S) or no evidence of growth (N; Fig 1). An extra category was added after the first farm visit 2 weeks after treatment as there were growths that could not be categorised as either H or S – a scorn (SC; Fig 1). A horn was defined as having normal growth without abnormalities. A scur was defined as any abnormal growth with a hardened (keratinised) surface that could be felt by hand in the horn bud area. A scorn was defined as a soft and fibrous (observed when cut) growth with a wide base and usually a rounded tip. N was recorded when the skin was smooth and there was no horn growth; this was considered a success. Horn buds were categorised into the four groups fortnightly for 2 months and then monthly for a further 3 months by a trained observer. The observer remained blind to the treatments each kid received. Once growth was observed, that treatment was considered unsuccessful and if both horn buds had evidence of growth, the animal was no longer monitored in subsequent checks. The probability of success, scurs, horns or scorns for each treatment at the final observation are presented.

**Fig 1.**
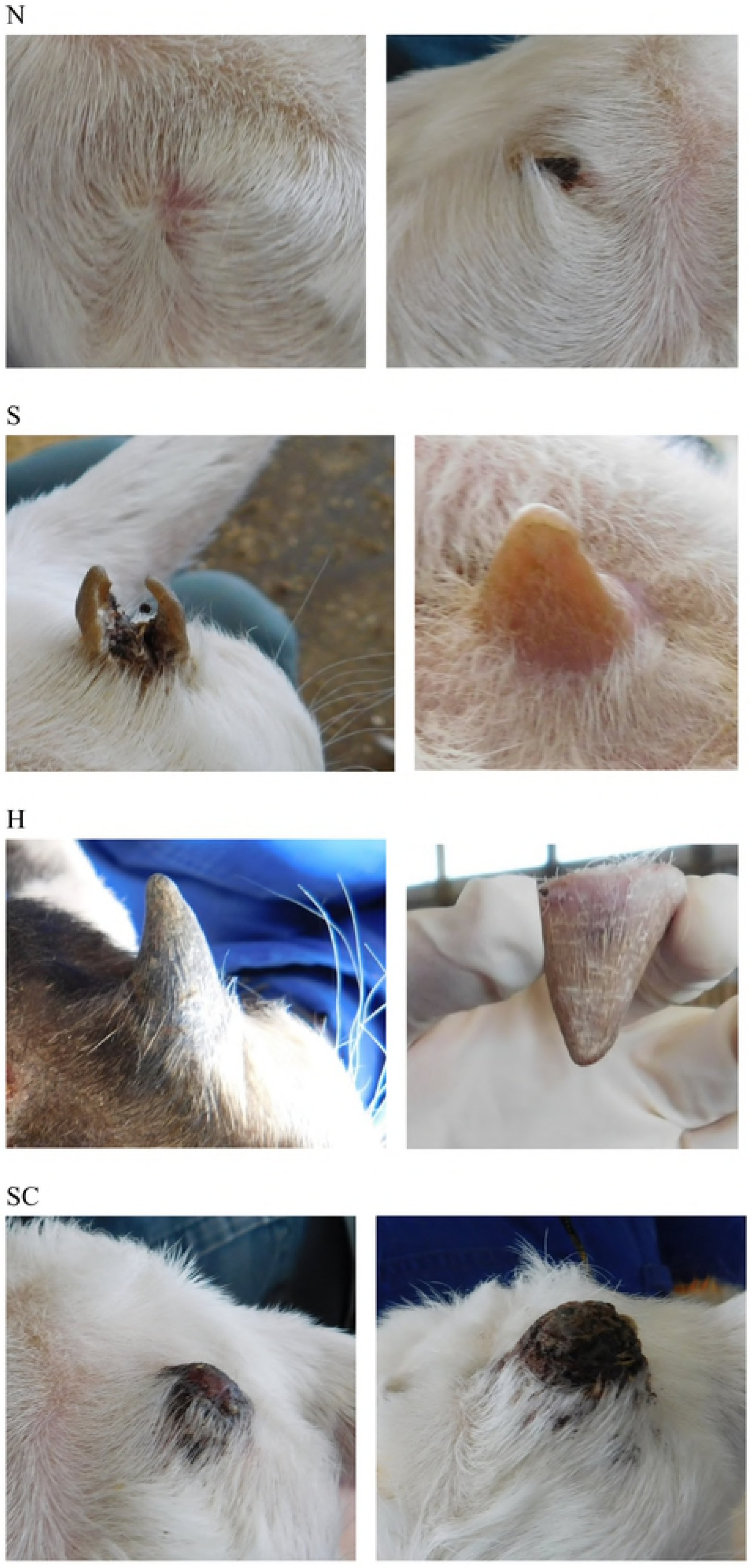
Categories of horn growth in dairy goat kids. No growth = N, scur = S, horn = H and scorn = SC. Goat kids were disbudded using either a clove oil injection (CLOVE), a cautery iron with the horn buds removed (BUDOFF) or a cautery iron with the horn buds left intact (BUDON).

## Statistical analysis

Genstat statistical software (version 18, VSN International, Hemel Hempstead, UK) was used to analyse the data. The binary response variables used for analyses included success (no growth = N), scurs (growth = S), horns (growth = H) and scorns (growth = SC); each response variable was assumed to be binomially distributed. Analysis of the differences between treatments were performed independently for all response variables. In addition, a bio-equivalence analysis was performed for the probability of success, with 80% power and an equivalence range of +/− 0.15. Bio-equivalence was assessed for each treatment with BUDOFF (considered the reference treatment) and also for CLOVE and BUDON (with BUDON as the reference treatment). We used a generalised linear mixed model for the analyses with a logit link. The fixed effects were for treatment and the treatment on the other horn bud (i.e. of the same kid). The random effects were for kid, farm, horn bud (left or right) and treatment date within farm. Differences between treatment means were compared using Fisher’s protected least significant difference test at the 5% significance level.

## Results

Of the 243 kids were enrolled in the study, 12 died before their 24 horn buds could be assessed 2 weeks following treatment; one animal (BUDON/BUDOFF) died 2 weeks post-treatment as a result of meningitis below the horn bud but the others died from complications not related to treatment (i.e. pneumonia, digestion issues). Data was missing for 9 BUDON and BUDOFF horn buds and 6 CLOVE horn buds. The differences in the probabilities of success, scurs, horns or scorns are presented in Fig 2. There was an effect of treatment on the probability of success (*F*_2,443_ = 43.3, *P* < 0.001). The probability of success for BUDOFF horn buds was higher than for BUDON and CLOVE horn buds (*P* ≤ 0.05). Furthermore, the probability of success for BUDON horn buds was higher than that of CLOVE horn buds (*P* ≤ 0.05). There was no evidence of bio-equivalence for the probability of success between BUDOFF and CLOVE, nor between BUDOFF and BUDON; however, the probabilities of success for BUDON and CLOVE horn buds were bio-equivalent.

**Fig 2.**
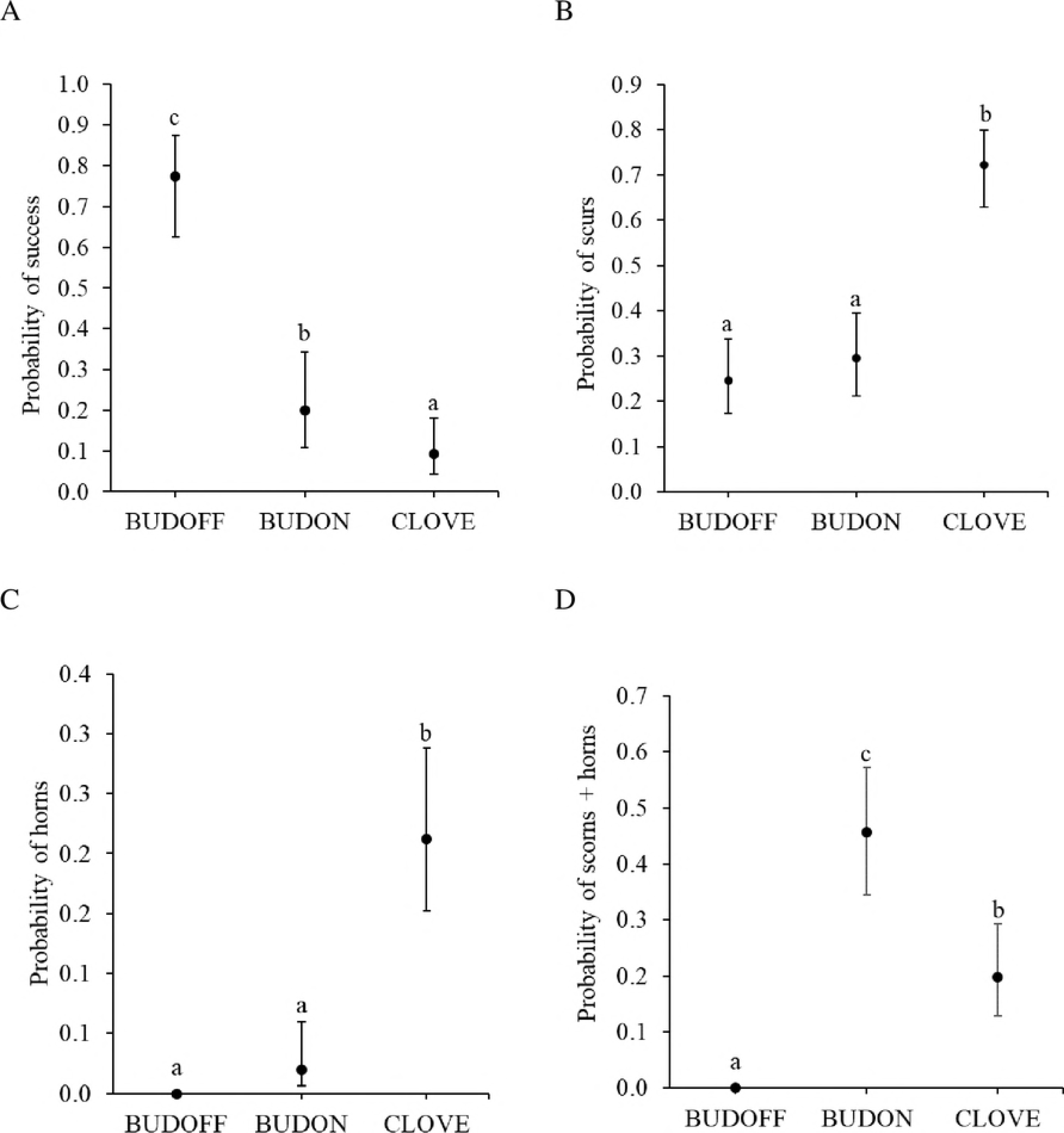
Probability of the four categories of horn growth with 95% confidence intervals for goat kids disbudded using three different techniques. (A) Success (no growth = N), (B) scurs (growth = S), (C) horns (growth = H) and (D) scorns and horn combined (growth = SC or H). Goat kids were disbudded using either clove oil injection (CLOVE; n = 156 horn buds), a cautery iron with the horn buds removed (BUDOFF; n = 153 horn buds) or a cautery iron with the horn buds left intact (BUDON; n = 153 horn buds). Means with differing subscripts are significantly different at *P* ≤ 0.05.

There was a treatment effect on the probability of developing scurs (*F*_2,452_ = 28.3, *P* ≤ 0.001). The probability of scurs was higher on CLOVE than BUDOFF and BUDON horn buds (P ≤ 0.05). There was no difference in the probability of scurs for BUDOFF and BUDON horn buds (P > 0.05).

There was a treatment effect for the probability of horns (*F*_2_ = 8.9 [chi-square test used as denominator degrees of freedom (ddf) were not estimable], *P* < 0.001). The probability of horns was higher for CLOVE than BUDON horn buds (*P* ≤ 0.05); horns were not observed for BUDOFF horn buds.

Scorns were only observed for BUDON horn buds (0.41 [0.25, 0.60]). There was an effect of treatment on the probability of horns and scorns combined (*F*_2_ = 11.2 [chi-square test used as ddf not estimable], *P* < 0.001). The probability of scorns and horns was higher for BUDON than CLOVE and BUDOFF horn buds (*P* ≤ 0.05).

## Discussion

The effectiveness of clove oil injection and two methods of cautery disbudding (i.e. horn buds removed vs. left intact) in preventing horn growth were evaluated in dairy goat kids. Clove oil injection has been previously reported by Molaei et al. [20], to be 100% successful in preventing horn growth in kids as well as calves [19]. In the present study, the CLOVE treatment appeared to be less effective at preventing horn growth than either of the cautery disbudding methods based on the high incidence of scurs and horns. Clove oil injection is a novel method of disbudding for goat kids compared with cautery disbudding (adapted from use in calves), which is the most commonly used method for disbudding goat kids worldwide [1]. Higher proportions of scurs on horn buds treated with clove oil compared with cautery disbudding may be associated with difficulties in restricting movement of the head during treatment. Clove oil was applied using a needle injected laterally into the buds whereas cautery disbudding involved pressing an ergonomic cautery iron down on the head. Consistent administration of the full volume of clove oil to the correct location (centre of the horn bud) was not always possible. The injection is likely to cause discomfort or pain [9] and kids generally struggled (rapid jerks of the head) during the procedure, resulting in the needle becoming dislodged. The creation of an applicator that can quickly deliver a consistent volume of clove oil to the right location may improve efficacy.

Potential explanations for the differences in efficacy of clove oil between Molaei et al. [20] and the present study include differences in methodologies. Molaei et al. [20] used a small sample size (16 vs. 243 kids in the present study), and their own clove oil distilled from the spice (clove oil used in the present study was sourced from a commercial manufacturer). The distilled clove oil appeared to be more effective as it totally prevented horn growth in both doe and buck kids which is notable as horn growth is more precocious in bucks [22]. The exact method of clove oil administration was not completely described in Molaei et al. [20] (e.g., it was not clear how the clove oil was injected or whether the hair was clipped so that any growth could be clearly observed). Perhaps more importantly, Molaei et al. [20] measured horn growth, whereas we evaluated any growth including horns, scurs (of any size) and scorns. It appears as though our methodology for injecting clove oil does not consistently prevent horns or scurs in goat kids and may not be a useful alternative to cautery disbudding with horn buds removed. Interestingly, there was a similar level of success in preventing scurs or horns between CLOVE and BUDON horn buds, indicating that leaving the horn bud intact may also not prevent horn regrowth. The method used in the present study to evaluate horn bud growth may be more comprehensive than methods used by farmers or farm staff and previous studies [19,20]. Future research should examine the exact mechanisms of clove oil action on the horn bud tissue of goat kids.

We observed some animals with inflammation of the upper eye lid area below the horn bud associated with the CLOVE treatment. This is interesting, as eugenol, the main component of clove oil, has anti-inflammatory properties [16,23]. Perhaps blood flow to the horn bud is reduced due to localised cellular necrosis and the observed swelling was pooling of blood below the horn bud. Measures of tissue sensitivity such as pressure algometry or Von Frey filaments, could be used to evaluate whether this apparent inflammatory response is painful.

We also had one kid (treated with BUDOFF and BUDON) die as a result of meningitis, which is likely associated with cautery iron use. It is well-established that cautery iron use can cause damage and thermal injury to the skull and brain, resulting in meningitis [11–13]. Although only one out of 243 kids died as a result of cautery disbudding, this demonstrates the capacity for cautery disbudding to not only cause pain, but mortalities.

To the authors’ knowledge, this is the first study to quantify the efficacy of cautery disbudding methods for goat kids. The BUDOFF treatment appeared to be most effective in preventing horns, scurs or scorns compared with the other two methods; this may be due to more complete horn bud destruction as the buds were cauterised and removed. By removing the horn bud, it is easier to ensure that all of the horn bud tissue is destroyed. It is generally considered more efficacious to remove the horn buds than leaving them on [1] and this method has been favoured in other studies [7,24,25].

The BUDON method resulted in abnormal growths that were soft, fibrous and fitted into neither the horn nor scur categories. The cautery iron used in the present study had a hollow centre with hot edges that cut into the skin, which may have allowed for the inner cells of the horn bud to continue to grow. This result has implications for farmers and contractors as it suggests that removing the horn buds may be more effective for preventing scurs than burning alone.

There is limited information available on horn growth in domesticated farmed goats, therefore comparisons with the literature are difficult. There is however, information on the horn growth of wild adult populations of goats [26], ibex [27] and chamois [28], usually with respect to environmental effects. Further research on scur or horn growth rates in goat kids after disbudding could help quantify the efficacy of cautery disbudding methods.

In order to understand regrowth associated with disbudding, it is important to understand how horns grow. The process involves keratinisation of the horn bud epidermis and ossification of the underlying dermis and hypodermis [29]. Goat horn anatomy is similar to that of cattle horns; the horn comprises tightly packed tubules produced by corium and germinal epithelium, which is attached to the frontal bone. The cornual diverticulum of the frontal sinus forms a cavity within the horn [1]. If the horn bud epidermis is not completely destroyed then keratinisation of some epidermal cells can occur resulting in scurs. Scurs usually result from inadequate burning of the horn bud site in an effort to reduce the risk of thermal injury to the skull and brain [22] or insufficient removal of the germinal tissue from the base of the horn bud [15]. If the base of the horn bud, which can be hard to see, is wider than the diameter of the cautery iron and not all tissue is destroyed, then scurs may also develop. Interestingly, in many cattle breeds, there is a gene for scurs [30–32], as well as polled and horned genes [33], meaning that scurs can grow naturally without disbudding. It is unknown whether a gene for scurs exists in goats. If so, herds with high rates of scurs may be associated with the genetic potential to grow scurs.

The best alternative to reduce pain and complications associated with cautery disbudding, would be to breed polled goats to eliminate the need for this procedure. There are hurdles to achieving this however, as the gene for polledness is linked with the intersex gene in goats [34]. This means that there is a high probability that naturally polled goats will be infertile. Until a polled line can be established, mitigating pain associated with cautery disbudding or further alternatives that cause less pain or injury than cautery disbudding should be considered.

## Conclusions

Clove oil injection did not appear to be as successful at preventing scurs or horns as cautery disbudding and therefore the methodology used in this study may not be useful for disbudding goat kids (if having no scurs is a concern). Interestingly cautery disbudding by removing the horn bud germinal tissue was more efficacious than leaving the horn buds intact, suggesting that the former method may be more effective for preventing horns, scurs or scorns. Future research should explore different clove oil formulations and administration methods or other alternatives to cautery disbudding that may be efficacious and cause less pain.

## Acknowledgements

We are grateful for assistance provided by AgResearch staff and students especially Ashleigh Weatherall, Rose Greenfield, Kirsty Owens, Rosalie Carter, Bridget Wise, Frances Huddart, Ariane Bright, Trevor Watson, Stephanie Delaney, Remi Mathieu, Gemma Lowe, Suzanne Dowling, Laura Hunter, Ben Ferniot and Johan Larive. We also thank Ali Cullum for her many contributions to this study and performing the disbudding treatments and the participating DGC farmers and farm staff. We also acknowledge assistance from Alan Julian for carrying out the pathology work. We are grateful to Cheryl O‘Connor and Vicki Burggraaf for reviewing this paper. This work was supported by the New Zealand Ministry of Business, Innovation and Employment (MBIE: Contract C10X1307; Wellington, New Zealand) and the Dairy Goat Cooperative (DGC) Ltd. (Hamilton, New Zealand).

